# Autophagy is impaired in fetal hypoplastic lungs and rescued by administration of amniotic fluid stem cell extracellular vesicles

**DOI:** 10.1101/2022.01.12.476078

**Authors:** Kasra Khalaj, Lina Antounians, Rebeca Lopes Figueira, Martin Post, Augusto Zani

## Abstract

**Rationale:** Pulmonary hypoplasia secondary to congenital diaphragmatic hernia (CDH) is characterized by reduced branching morphogenesis, which is responsible for poor clinical outcomes. Administration of amniotic fluid stem cell extracellular vesicles (AFSC-EVs) rescues branching morphogenesis in rodent fetal models of pulmonary hypoplasia. Herein, we hypothesized that AFSC-EVs exert their regenerative potential by affecting autophagy, a process required for normal lung development.

**Objectives:** To evaluate autophagy in hypoplastic lungs throughout gestation and establish whether AFSC-EV administration improves branching morphogenesis through autophagy-mediated mechanisms.

**Methods:** EVs were isolated from c-kit+ AFSC conditioned medium by ultracentrifugation and characterized for size, morphology, and EV markers. Branching morphogenesis was inhibited in rat fetuses by nitrofen administration to dams and in human fetal lung explants by blocking RAC1 activity with NSC23766. Expression of autophagy activators (BECN1 and ATG5) and adaptor (SQSTM1/p62) was analyzed *in vitro* (rat and human fetal lung explants) and *in vivo* (rat fetal lungs). Mechanistic studies on rat fetal primary lung epithelial cells were conducted using inhibitors for microRNA-17 and -20a contained in the AFSC-EV cargo and known to regulate autophagy.

**Measurements and Main Results:** Rat and human models of fetal pulmonary hypoplasia showed reduced autophagy mainly at pseudoglandular and canalicular stages. AFSC-EV administration restored autophagy in both pulmonary hypoplasia models by transferring miR-17∼92 cluster members contained in the EV cargo.

**Conclusions:** AFSC-EV treatment rescues branching morphogenesis partly by restoring autophagy through miRNA cargo transfer. This study enhances our understanding of pulmonary hypoplasia pathogenesis and creates new opportunities for fetal therapeutic intervention in CDH babies.

## Introduction

Fetal lung underdevelopment, also known as pulmonary hypoplasia, is characterized by reduced branching morphogenesis (1). The most common cause of pulmonary hypoplasia is congenital diaphragmatic hernia (CDH), a birth defect due to an incomplete closure of the fetal diaphragm and herniation of abdominal organs into the fetal chest. Despite decades of research, CDH continues to have high morbidity and mortality rates (2), which are directly related to the severity of lung underdevelopment (3). There is consensus that intervening after birth is too late, as some fetuses die *in utero* or are terminated, some succumb after birth, and those who survive and undergo surgery never regain normal lung growth (4).

We have recently demonstrated that administration of extracellular vesicles derived from amniotic fluid stem cells (AFSC-EVs) enhances fetal lung development by promoting branching morphogenesis and rescuing epithelial cell differentiation and homeostasis in various fetal rodent models of pulmonary hypoplasia (5). EVs are membrane-bound nanoparticles that mediate intercellular communication and are responsible for stem cell paracrine signaling (6). EVs carry cargo in the form of bioactive molecules, including genetic material (7–9). Investigating AFSC-EV mechanism of action, we found that their beneficial effects on fetal lung tissue were exerted, at least in part, via miRNAs that regulate the expression of genes involved in lung development (5). Particularly, AFSC-EVs were highly enriched in members of the miR17∼92 cluster, which is known to control lung branching morphogenesis (9), lung progenitor cell proliferation and differentiation (10, 11), surfactant protein C secretion (12), and alveolarization (11). However, the mechanisms underlying the lung regenerative effects of AFSC-EV administration remain to be fully elucidated.

Autophagy has recently been reported as one of the biological processes required to coordinate normal lung development (13). Specifically, impaired autophagy compromises lung branching morphogenesis, thereby contributing to reduced saccular formation in bronchopulmonary dysplasia (BPD), a condition characterized by lung immaturity secondary to premature birth (13). Autophagy is an evolutionary conserved process that provides protective functions to damaged cells and organs (14). It is considered essential for the maintenance of cellular homeostasis in the airway epithelium (15), and several studies have shown that impaired autophagy leads to epithelial cell dysfunction (16). As the lungs of BPD patients are immature similar to those of babies with pulmonary hypoplasia secondary to CDH, we investigated whether autophagy is dysregulated also in fetal hypoplastic lungs.

Herein, we report that autophagy signaling is impaired in rodent and human models of fetal pulmonary hypoplasia at different stages of lung development. Moreover, we demonstrate that administration of AFSC-EVs rescues branching morphogenesis in both species in part through restoration of autophagy via miR-17∼92 cargo transfer. This study increases our understanding of the biological mechanisms contributing to pulmonary hypoplasia and creates new opportunities for fetal therapeutic intervention.

## Materials and Methods

### Extracellular Vesicles

EVs were derived from c-kit+ rat AFSCs and good manufacturing practices (GMP)-grade human AFSCs using our published protocol (5, 17). Following the International Society for Extracellular Vesicles guidelines, EVs were characterized for size (nanoparticle tracking analysis), morphology (transmission electron microscopy), and expression of canonical EV-related protein markers (Western blot analysis) (**Figure E1**), as described (5, 18).

### Experimental models of pulmonary hypoplasia

Following ethical approval of the Animal Care Committee at the Hospital for Sick Children (AUP#49892), embryonic (E) day 9.5 dams were gavaged with either nitrofen to induce fetal pulmonary hypoplasia or olive oil (vehicle) as control, as described (19, 20). For rat explant studies, lungs from nitrofen-exposed and control pups were micro-dissected at embryonic (E12.5), pseudoglandular (E14.5), canalicular (E17.5), and saccular (E20.5) stages, grown on nanopore membranes, and incubated for 72 hours in culture medium alone or with rat AFSC-EVs (0.5% v/v), as described (5).

For primary cell studies, single cell suspensions of E14.5 and E17.5 lungs were enriched for epithelial cells using an established protocol (5, 21, 22).

For *in vivo* administration of AFSC-EVs, we anaesthetized rat dams at E18.5, exposed uterine horns through midline laparotomy, and injected amniotic sacs with either 100 μL of saline or rat AFSC-EVs (6×10^8^ ± 1×10^6^ vesicles). We returned the uterine horns into the abdomen, closed the laparotomy, and harvested the fetal lungs after 72 hours.

Human fetal lungs were obtained under informed consent from Mount Sinai Hospital Biobank, Toronto (REB#10-0128-E), from 8 healthy fetuses terminated at a median of 17 weeks of gestation (range, 16-19), and grown as explants using an adapted protocol (23). Briefly, 1-2 mm^3^ pieces were dissected from the left lower lobe and grown on nucleopore membranes for 96 hours. To inhibit branching morphogenesis, we administered NSC23766 (25 μM, refreshed every 24 hours for a total of 48 hours) as described (23). NSC23766 is an inhibitor of RAC1, a Rho GTPase that modulates lung branching morphogenesis and whose expression is downregulated in lungs of CDH fetuses, as confirmed with immunofluorescence staining on autopsy samples (REB#1000074888) (**Figure E2**). Injured explants were treated with medium alone or medium supplemented with human AFSC-EVs (0.5% v/v) at 48 and 72 hours. Uninjured (no NSC23766), untreated (no AFSC-EVs) explants were used as control.

### Branching morphogenesis and autophagy evaluation

To assess lung branching morphogenesis, rat fetal lung explants were imaged by differential interference contrast microscopy (embryonic/pseudoglandular stages) or processed for histology (hematoxylin/eosin staining for canalicular/saccular stages) as described (5). Explants were blindly evaluated for terminal bud density at embryonic/pseudoglandular stages as described (5), and for airspace density using the radial airspace count (RAC) at canalicular/saccular stages as recommended by the American Thoracic Society (24).

To study autophagy, we determined gene and protein expression of autophagy activators (Beclin 1, BECN1; Autophagy Related 5, ATG5) and adaptor (Sequestosome 1, SQSTM1 also known as p62). For gene expression, we conducted RNA extraction, cDNA synthesis and qPCR quantification as described (5, 25). Primers are listed in **Table 1**. For protein expression, we performed Western blot and immunofluorescence assays, as described (5). Antibodies and concentrations are specified in **Table 2**. To evaluate autophagosome density, we used transmission electron microscopy as described (13).

**Table 1.**
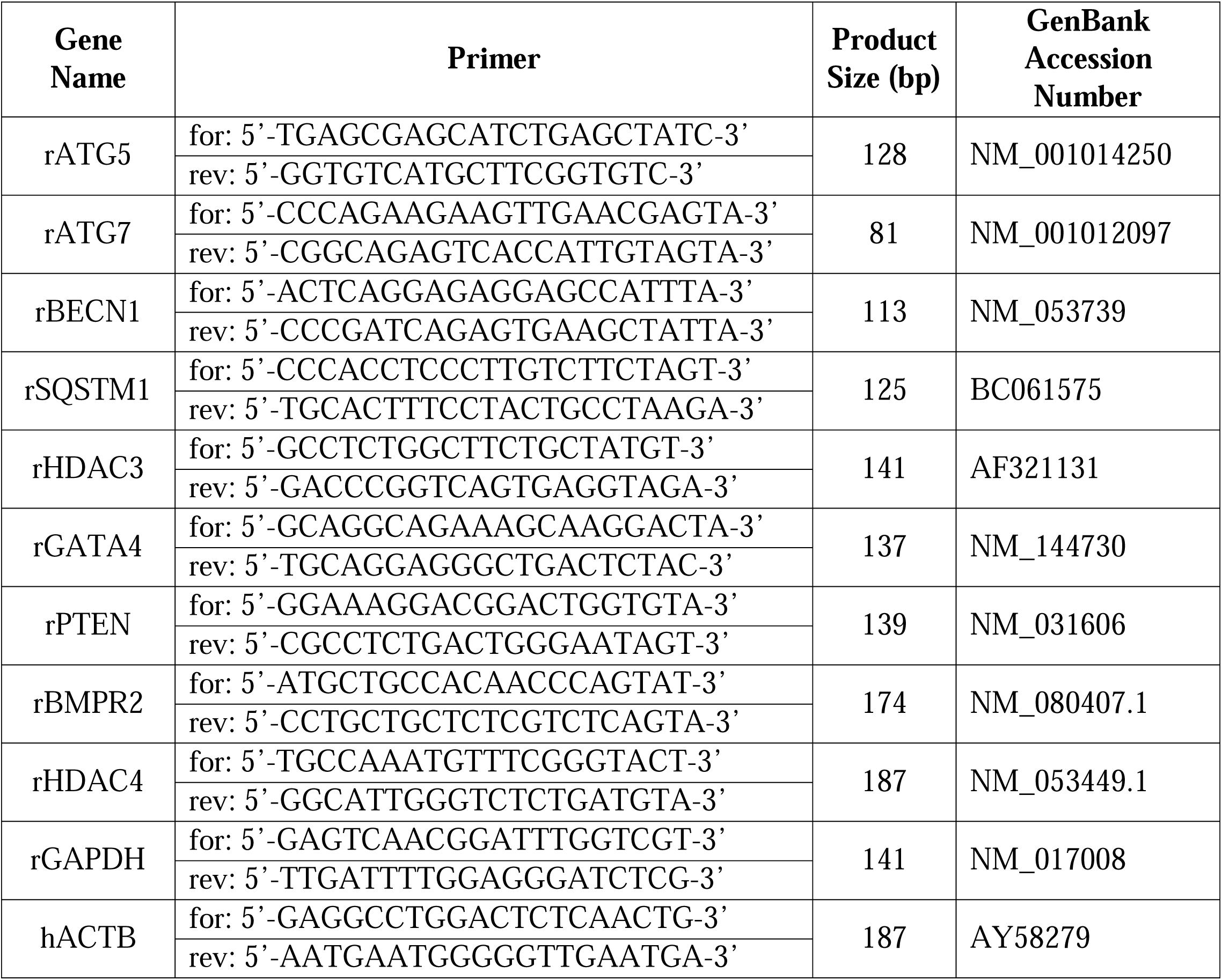
mRNAs assessed by Real-Time PCR.

**Table 2.** Antibodies utilized for imaging and Western blotting.

| Gene name | Antibody | Antibody host | Manufacturer/Catalogue # | Application | Dilution Factor |
| --- | --- | --- | --- | --- | --- |
| Beclin-1 | BECN1 | Mouse | St John's/SJT97770 | WB, IF | 1:500 (IF),<br>1:1000 (WB) |
| Sequestosome-1/p62 | SQSTM1/p62 | Rabbit | CST/5114 | WB, IF | 1:300 (IF),<br>1:1000 (WB) |
| Epithelial cell adhesion molecule | EPCAM/CD236 | Rabbit | Abcam/ab213500 | IF | 1:250 |
|  | Epcam/CD236 | Mouse | eBioscience/14-5791 | IF | 1:300 |
| Vimentin | Vimentin | Rabbit | CST/3932 | IF | 1:500 |
| Autophagy related 5 | ATG5 | Rabbit | CST/2630S | WB | 1:1000 |
| GATA binding protein 4 | GATA4 | Rabbit | Abcam/ab84593 | WB | 1:1000 |
| Phosphatase and tensin homolog deleted on chromosome 10 | P-PTEN | Rabbit | Sigma/SAB4300044 | WB | 1:1000 |
| Histone deacetylase 3 | HDAC3 | Mouse | CST/3949 | WB | 1:1000 |
| Microtubule-associated proteins 1A/1B light chain 3B | LC3B | Rabbit | CST/2775S | WB | 1:1000 |
| Ras-related C3 botulinum toxin substrate 1 | RAC1 | Mouse | Abcam/ab33186 | IF | 1:100 |
| Tumor susceptibility 101 | TSG101 | Mouse | Santa-Cruz/Sc79644 | WB | 1:1000 |
| Flotillin-1 | Flotillin-1 | Mouse | BD/610820 | WB | 1:500 |
| CD63 | CD63 | Mouse | Abcam/ab59479 | WB | 1:500 |
| Calnexin | Calnexin | Rabbit | Abcam/ab22595 | WB | 1:1000 |
| Not applicable | IgG Alexafluor-488 | Rabbit | CST/4412S | IF | 1:1000 |
|  | IgG Alexafluor-555 | Mouse | CST/4409 | IF | 1:1000 |
|  | IgG HRP | Mouse | CST/7076S | WB | 1:1000 |
|  | IgG HRP | Rabbit | CST/7074 | WB | 1:1000 |

CST: Cell signaling technology
WB: Western blot
IF: Immunofluorescence

### miRNA inhibition studies

To inhibit miRNAs of interest, we treated rat AFSCs at 60% confluence with miR-17-5p and - 20a power inhibitors (25 μM) following the manufacturer’s protocol (Qiagen YI04100215-DDA; miR-17-5p, YI04100414-DDA; miR-20a, YI00199006-DDA; Scrambled Negative Control). Briefly, cells were subjected to either inhibitor or a negative control for 36 hours. EVs were harvested from cultured medium, as described (5). miRNA inhibition was validated by assessing the gene expression of known miR-17∼92 targets (BMPR2; Bone morphogenetic protein receptor type II, HDAC4; Histone deacetylase 4, ATG7; Autophagy related 7) (26–28) (**Figure E3**). Nitrofen exposed primary lung epithelial cells isolated at E17.5 were treated with the modified EVs and assessed for expression of autophagy markers. Fetal lung explants treated with the modified EVs were analyzed for airspace density using radial airspace count (RAC).

### Statistical analysis

Groups were compared using two-tailed Student t-test, Mann-Whitney test, one-way analysis of variance (Tukey’s post-test), or Kruskal-Wallis test, as appropriate. P < 0.05 was considered significant.

## Results

### Autophagy is impaired in rat fetal hypoplastic lungs at different developmental stages

We first confirmed that administration of nitrofen to rat dams at E9.5 reduced branching morphogenesis in the lungs of fetuses during all four stages of fetal lung development (**Figure 1A-D**). Specifically, compared to fetuses not exposed to nitrofen, lung explants harvested from nitrofen-exposed rat pups had fewer buds at the embryonic (E12.5), pseudoglandular (E14.5), canalicular (E17.5), and saccular (E20.5) stages. When we analyzed the gene expression of critical autophagy markers, we found that autophagy activators BECN1 and ATG5 were downregulated in nitrofen-exposed lungs (**Figure 1E-F**). Moreover, expression levels of SQSTM1, an autophagy adaptor whose high levels are indicative of autophagy disruption (29), were upregulated in nitrofen-exposed lungs compared to control (**Figure 1G**). Since all three autophagy markers were dysregulated at the pseudoglandular and canalicular stages, we focused on these two lung developmental stages and confirmed the dysregulation by immunostaining (**Figure 1H-K**) and transmission electron microscopy (**Figure 1L-M**). Particularly, compared to control, hypoplastic lung explants had fewer autophagosomes, which are active autophagy units, at both stages (**Figure 1L-M**).

**Figure 1.**
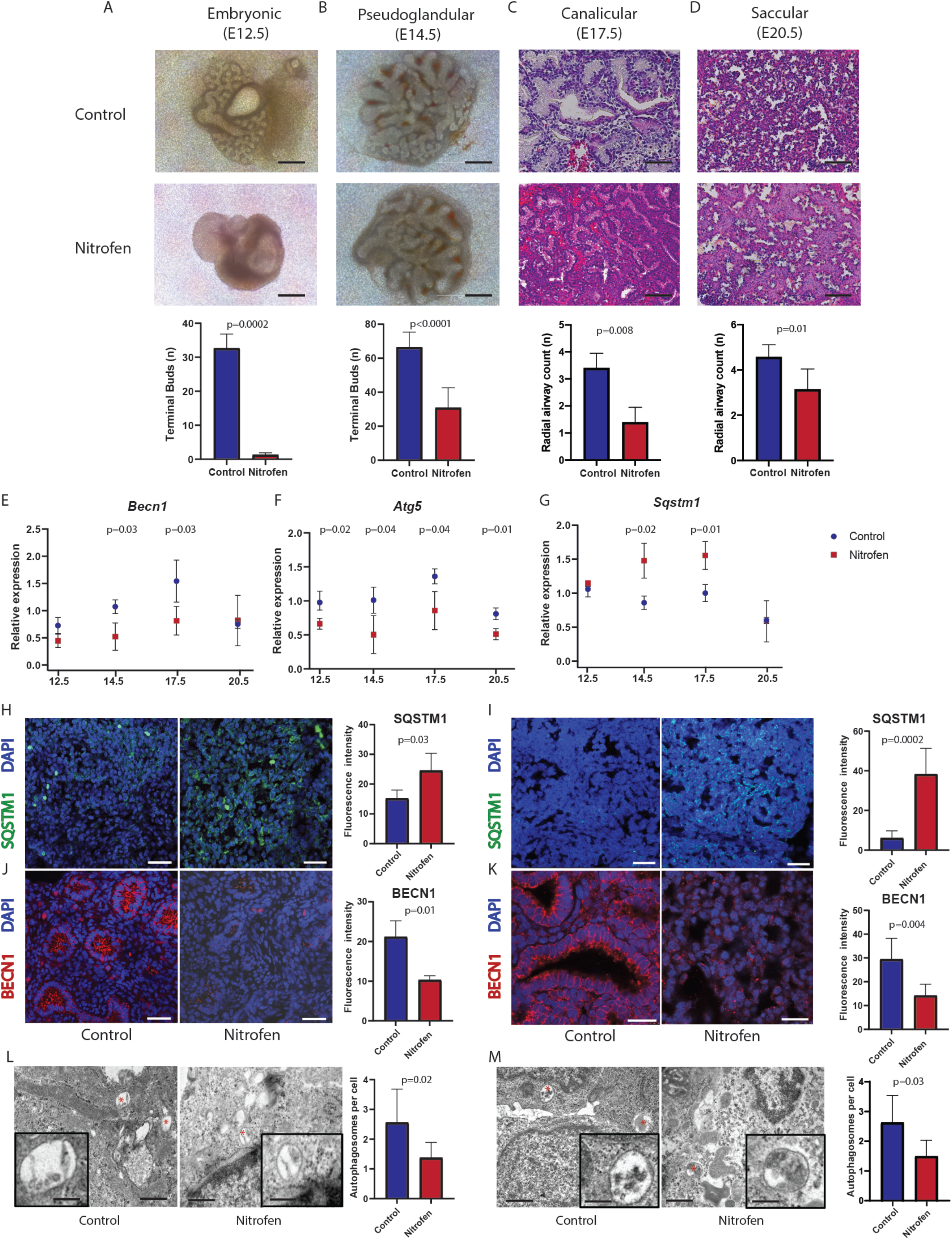
Branching morphogenesis and autophagy are impaired during fetal lung development in an experimental rat model of congenital diaphragmatic hernia (CDH). (*A-D*) Nitrofen-exposed fetal lung explants had reduced branching morphogenesis in all four stages of fetal lung development. Branching morphogenesis was evaluated as number of terminal buds at embryonic and pseudoglandular stages and as airspace density using radial airspace count at canalicular and saccular stages. *N*= 6 pups per group. Scale bars, 50 μm (*A-B*) and 100 μm (*C-D*). (*E-G*) Time course analysis of autophagy marker BECN1, ATG5, and SQSTM1 gene expression during fetal lung development showed highest autophagy impairment at the pseudoglandular and canalicular stages. *N*= 3 pups per group. (*H-K*) Representative immunofluorescence and quantification of BECN1 and SQSTM1 expression in fetal lung explants of control and nitrofen-exposed rats. *N*= 6 pups per group (9 fields per animal). Scale bars, 25 μm. (*L-M*) Representative transmission electron microscopy images and quantification of autophagosomes (red asterisks) in fetal lung explants showing lower density in nitrofen-exposed rats. Inlets depict higher magnification of representative autophagy units. *N*= 9 cells per group (9 fields per animal, 3 animals per condition). Scale bars: 1 μm (inlet scale bars: 100 nm). For all panels, data are shown as mean ± SD and were analyzed using Student’s t-test (*A-B, E-M*) or Mann-Whitney test (*C-D*). Only significant differences (p<0.05) are reported in the graphs.

### Administration of AFSC-EVs restores impaired autophagy in rat fetal hypoplastic lungs

We first sought to investigate the effects of rat AFSC-EV administration on the branching morphogenesis of rat fetal hypoplastic lungs. We confirmed that AFSC-EV-treated hypoplastic lung explants harvested during the pseudoglandular (E14.5) and canalicular (17.5) stages had an increased number of airspaces (**Figure E4**). When we assessed the effects of rat AFSC-EVs on autophagy at the pseudoglandular and canalicular stages, we observed restoration of BECN1, ATG5, and SQSTM1 gene expression (**Figure 2A-B**). Moreover, AFSC-EV treated lung explants had rescued BECN1 and SQSTM1 protein expression at the pseudoglandular stage (**Figure 2C**), and BECN1, ATG5, and SQSTM1 protein expression at the canalicular stage (**Figure 2D**). We also observed that in AFSC-EV treated lung explants, the density of autophagosomes per cell was restored to control levels at both stages (**Figure 2E-F**).

**Figure 2.**
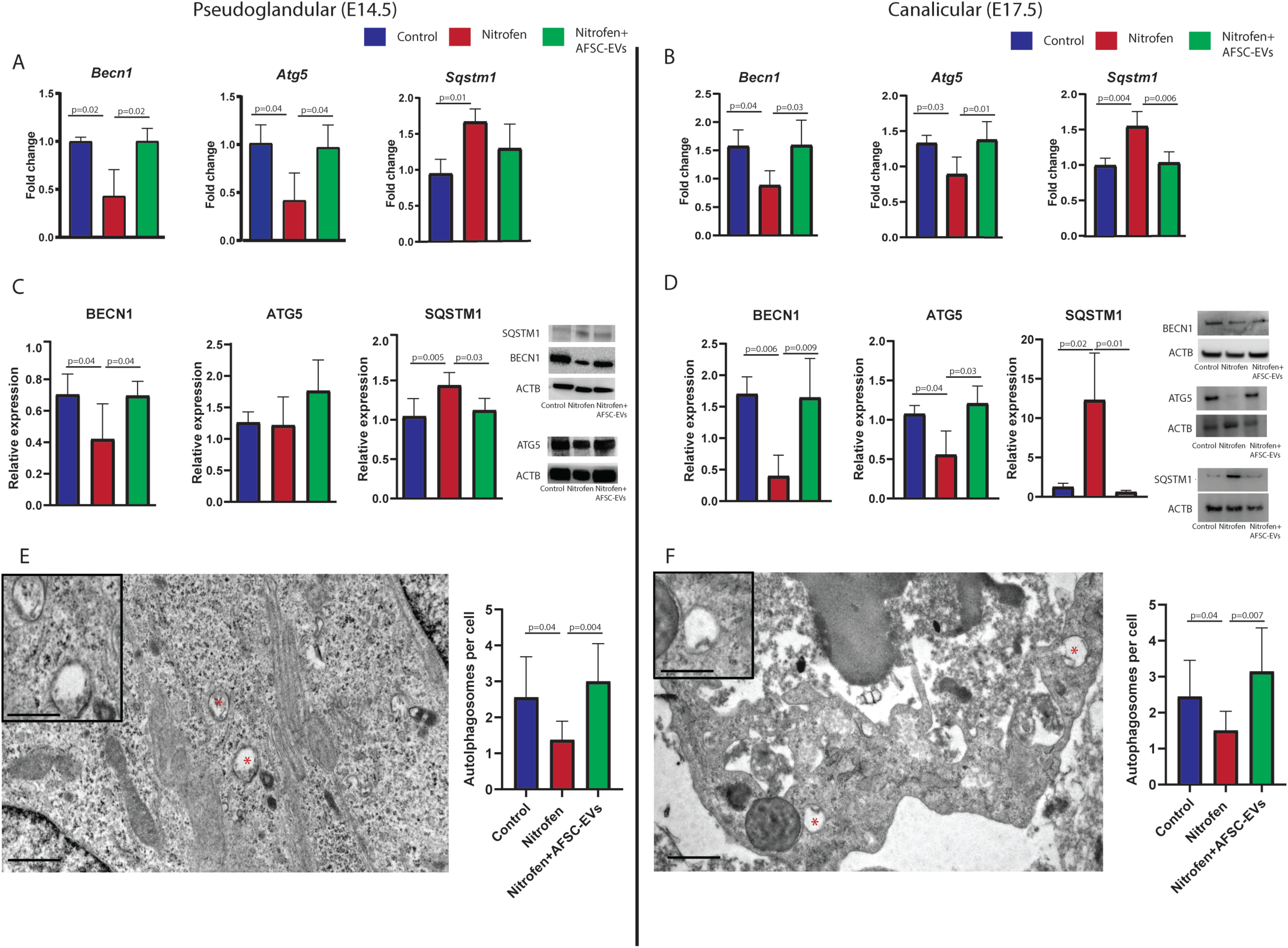
Impaired autophagy in fetal hypoplastic lungs is rescued by administration of amniotic fluid stem cell extracellular vesicles (AFSC-EVs). (*A-D*) Autophagy markers measured by gene and protein expression were impaired in nitrofen-exposed lungs and rescued by AFSC-EV administration. *N*=4 pups per group. (*E-F*) Representative transmission electron microscopy images and quantification of autophagosomes (red asterisks) indicating that AFSC-EV administration restored autophagosome density. Inlets depict higher magnification of representative autophagy units. *N*= 9 cells per field per group, 3 animals per condition. Scale bars: 1 μm (inlet scale bars: 100 nm). For all panels, data are shown as mean ± SD and were analyzed using One-way ANOVA with Tukey’s multiple comparison test. Only significant differences (p<0.05) are reported in the graphs.

### Impaired fetal lung autophagy is restored by members of the miR-17∼92 cluster contained in AFSC-EVs

As autophagy has been reported to be primarily activated in the epithelium of the mouse fetal lung (13), we studied the colocalization of BECN1 with EPCAM, an epithelial cell marker, in our rat fetal model. At immunofluorescence, we found that BECN1 was expressed in the cytoplasm of epithelial cells of all three conditions (**Figure 3A**). This confirmed autophagy activation in the epithelium, whose homeostasis is known to be dysregulated in hypoplastic lungs and rescued by the administration of AFSC-EVs (5, 22, 30). In our previous study, we showed that primary lung epithelial cells isolated from rat fetal hypoplastic lungs and treated with AFSC-EVs had higher levels of members of the miR-17∼92 cluster and their paralogues compared to untreated epithelial cells (5). When we previously correlated the AFSC-EV miRNA cargo content with the validated mRNAs in the target epithelial cells, we found that miR-17 and miR-20a negatively correlated with SQSTM1 mRNA (5) (**Figure E5**). In the present study, we confirmed that the effects of AFSC-EVs on autophagy in hypoplastic lungs were related to the miR-17∼92 cluster by conducting miRNA inhibition studies using miR-17 and -20a antagomirs (**Figure 3B**). We selected these two miRNAs, as they were the two most abundant members of the miR-17∼92 cluster in our AFSC-EV cargo sequencing analysis (5). We first transfected parental amniotic fluid stem cells (AFSCs) with inhibitors of either miR-17 or miR-20a, or with a control scrambled sequence (**Figure 3B**). We demonstrated that knockdown of both miRNAs resulted in upregulation of common target genes (BMPR2, HDAC4 and ATG7) in both parental AFSCs and in their secreted AFSC-EVs, thus validating miRNA inhibition (**Figure E3**). When treating fetal lung explants at E17.5, knockdown of both miRNAs caused a reduction of airspace density (**Figure 3C**). Inhibition of miR-20a in AFSC-EVs resulted in downregulation of BECN1 and upregulation of SQSTM1 gene expression in target primary lung epithelial cells isolated at E17.5 compared to cells treated with the scrambled sequence (**Figure 3D**). Similarly, inhibition of miR-17 upregulated SQSTM1 gene expression in target primary lung epithelial cells (**Figure 3D**). At the protein level, we observed that inhibition of both miR-17 and -20a resulted in downregulation of BECN1 and SQSTM1 (**Figure 3E**). We further corroborated these findings with the evidence of downregulation of LC3B-II, the activated autophagy complex isoform (**Figure 3E**).

**Figure 3.**
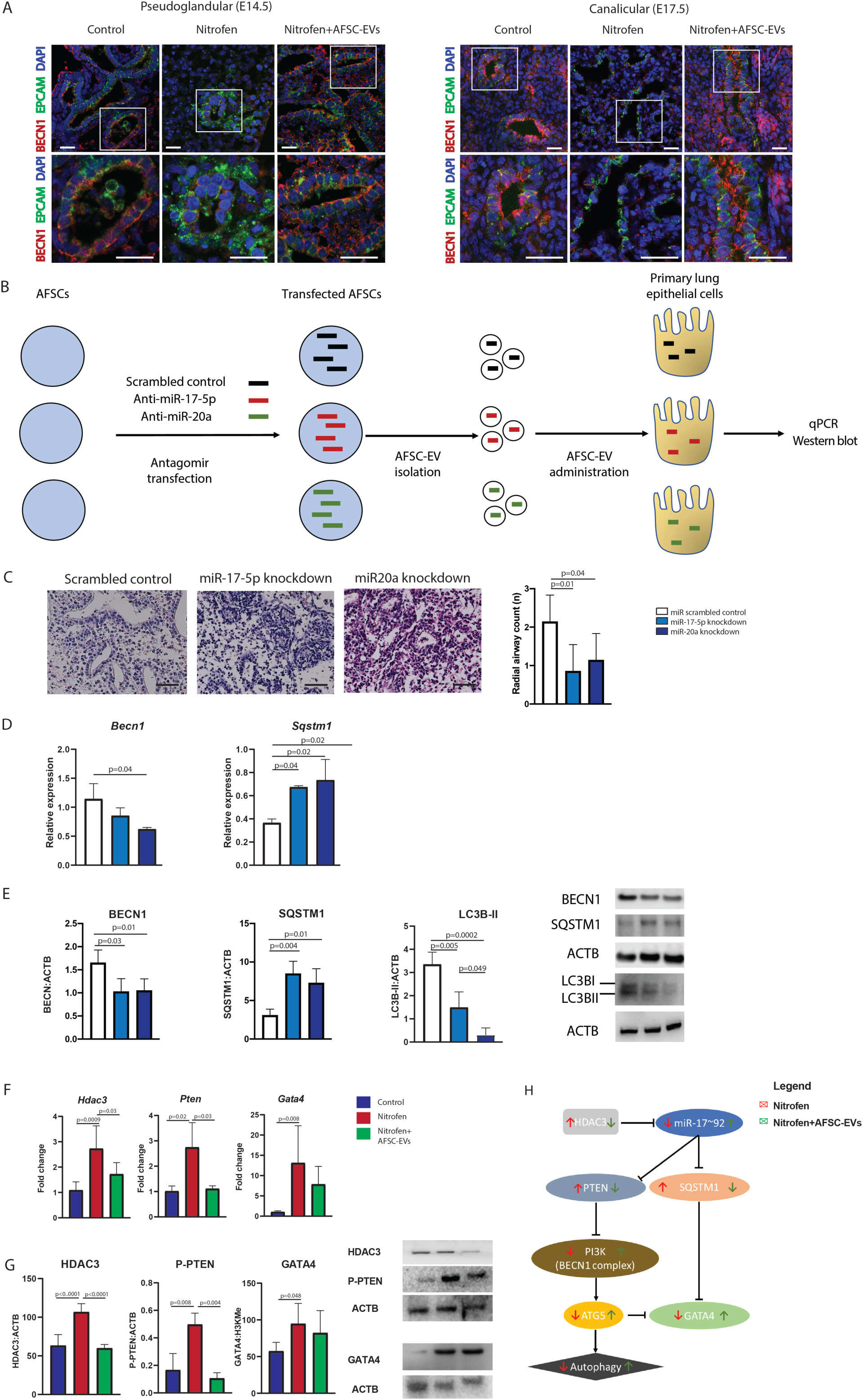
AFSC-EVs restore autophagy in the epithelium of fetal hypoplastic lungs via transfer of miR-17∼92 members contained in the AFSC-EV cargo. (*A*) Representative immunofluorescence images of the colocalization of BECN1+ cells with EPCAM+ cell showing autophagy activation in the lung epithelium in all three groups. *N*=6 pups per group. Scale bars, 25 μm, 75 μm (inlets). (*B*) Schematic of miRNA mechanistic studies. (*C*) Administration of modified AFSC-EVs to fetal rat lung explants at the canalicular stage results in decreased airspace density using radial airspace count. *N* ≥ 3 pups per group. Scale bars, 100 μm. *(D-E)* Gene and protein expression levels of autophagy markers in nitrofen-exposed primary lung epithelial cells treated with EVs derived from AFSCs treated with miR-17-5p and -20a antagomirs or scrambled sequence. *N*= 4 biological replicates per group. (*F-G*) Gene and protein expression levels of factors involved in the autophagy signalling pathway (HDAC3, PTEN, and GATA4) were upregulated in hypoplastic fetal lungs and downregulated by AFSC-EV treatment. *N*=6 pups per group. (*H*) Summary of the miR17∼92-autophagy axis with the effects of nitrofen injury (red arrows) and AFSC-EV treatment (green arrows). Data are shown as means ± SD and were analyzed using one-way ANOVA with Tukey’s multiple comparison test. Only significant differences (p<0.05) are reported in the graphs.

To further elucidate the effects of the AFSC-EV miR-17∼92 cluster on autophagy in our pulmonary hypoplasia model, we analyzed the expression of factors that are known to be regulated by the cluster and implicated in the autophagy signalling cascade. We first tested the expression of HDAC3, a factor that plays an important role in lung epithelial cell remodeling during the late stages of lung development via miR-17∼92 inhibition (31) and in autophagy regulation (32, 33). We found that in hypoplastic fetal lungs at the canalicular stage, HDAC3 expression was increased compared to normal lungs and rescued to normal levels by AFSC-EV administration (**Figure 3F-G**). When we investigated the expression of PTEN and GATA4, which are downstream factors in the autophagy pathway and directly or indirectly regulated by the miR-17∼92 cluster (34, 35), we found that hypoplastic lungs had higher levels than control (**Figure 3F-G**). Treatment with AFSC-EVs restored PTEN expression back to control levels but had no restorative effect on GATA4 expression (**Figure 3F-G**). The putative miR17∼92 autophagy axis in rat fetal lungs during lung development is schematically depicted in **Figure 3H**.

### Intra-amniotic injection of AFSC-EVs rescues autophagy in an *in vivo* fetal rat model of pulmonary hypoplasia

Towards the clinical translation of AFSC-EVs as a potential therapy for fetal lung underdevelopment, we investigated the effects of intra-amniotically administered rat AFSC-EVs on autophagy in E18.5 rat fetuses. We chose this gestational age as it corresponds to the timepoint when clustered rhythmic fetal breathing movements are first observed (36). Lungs were harvested at E21.5 and those exposed to nitrofen were hypoplastic as evidenced by decreased airspace density compared to control (**Figure 4A**). Lungs from fetuses that received an intra-amniotic injection of AFSC-EVs had rescued airspace density (**Figure 4A**). Nitrofen-exposed lungs that received an intra-amniotic injection of PBS had impaired autophagy, with lower BECN1 and ATG5 and higher SQSTM1 gene and protein expression compared to control (**Figure 4B**). Conversely, fetuses that received intra-amniotic AFSC-EV treatment had rescued BECN1, ATG5, and SQSTM1 gene and protein expression levels back to control (**Figure 4B-C**).

**Figure 4.**
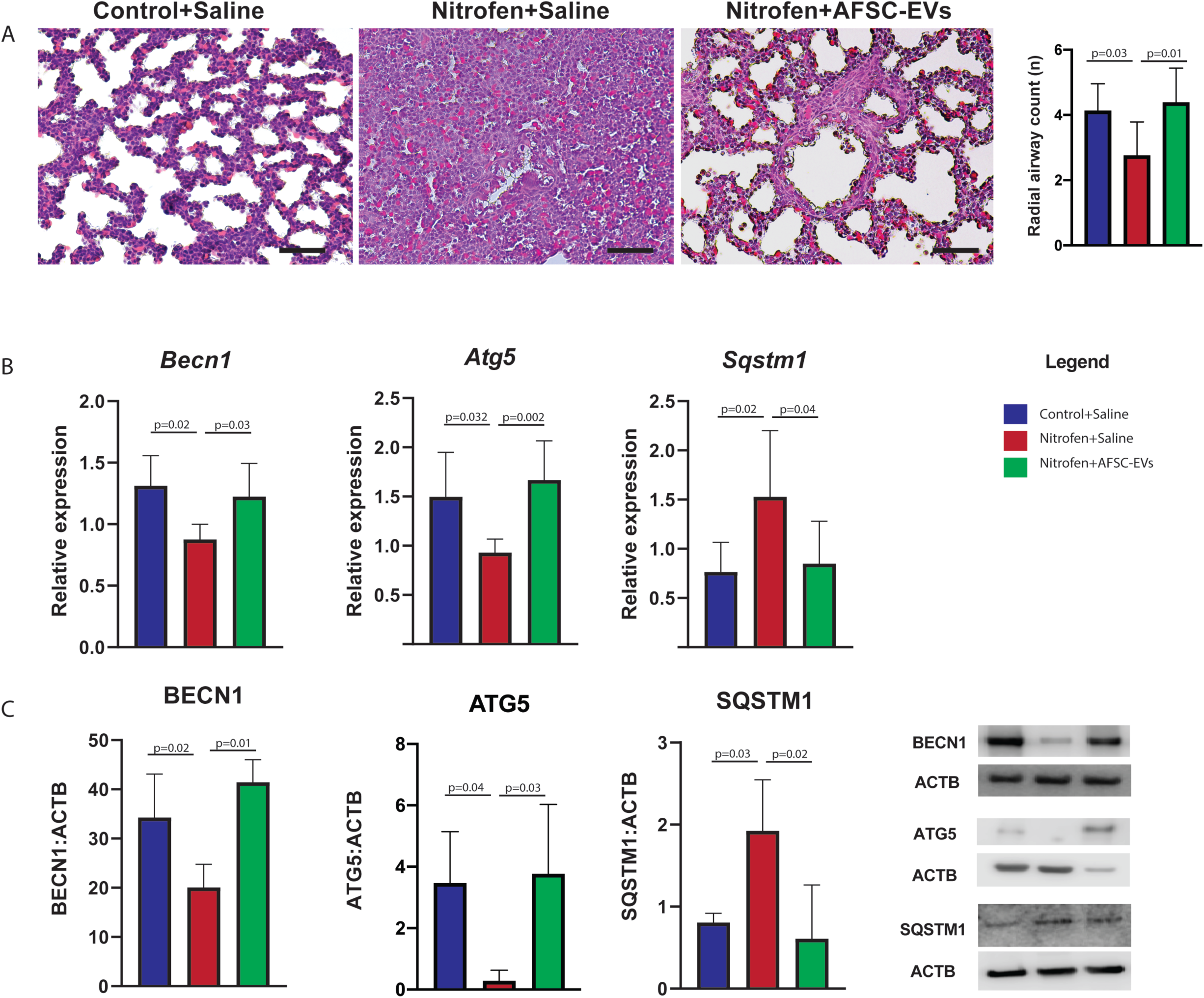
Intra-amniotic injection of AFSC-EVs rescues autophagy in an *in vivo* rat model of pulmonary hypoplasia secondary to CDH. (*A*) Representative histology images of fetal lung architecture and airspace density quantification using radial airspace count. Branching morphogenesis was reduced in nitrofen-exposed pups treated with saline and increased in fetuses treated with AFSC-EVs. Scale bars, 100 μm. *N*= 8 for each group. (*B-C*) Intra-amniotic treatment with AFSC-EVs resulted in restoration of autophagy gene and protein expression profiles. *N*≥ 5 per group. Data are shown as mean ± SD and were analyzed using One-way ANOVA with Tukey’s multiple comparison test. Only significant differences (p<0.05) are reported in the graphs.

### Impairment autophagy is recapitulated in an *ex-vivo* human model of pulmonary hypoplasia and restored by treatment with human AFSC-EVs

We next employed an *ex-vivo* human model of pulmonary hypoplasia using NSC23766, a RAC-1 inhibitor that inhibits branching morphogenesis as previously shown (23) (**Figure 5A**). BECN1 and ATG5 were downregulated in hypoplastic human fetal lungs and rescued to control levels with the administration of human GMP-grade AFSC-EVs (**Figure 5B-C**). Conversely, we did not find a difference in SQSTM1 expression between normal and hypoplastic lungs (**Figure 5B-C**). The density of autophagosomes was decreased in human fetal hypoplastic lungs compared to control and rescued by the administration of human AFSC-EVs back to control levels (**Figure 5D**).

**Figure 5.**
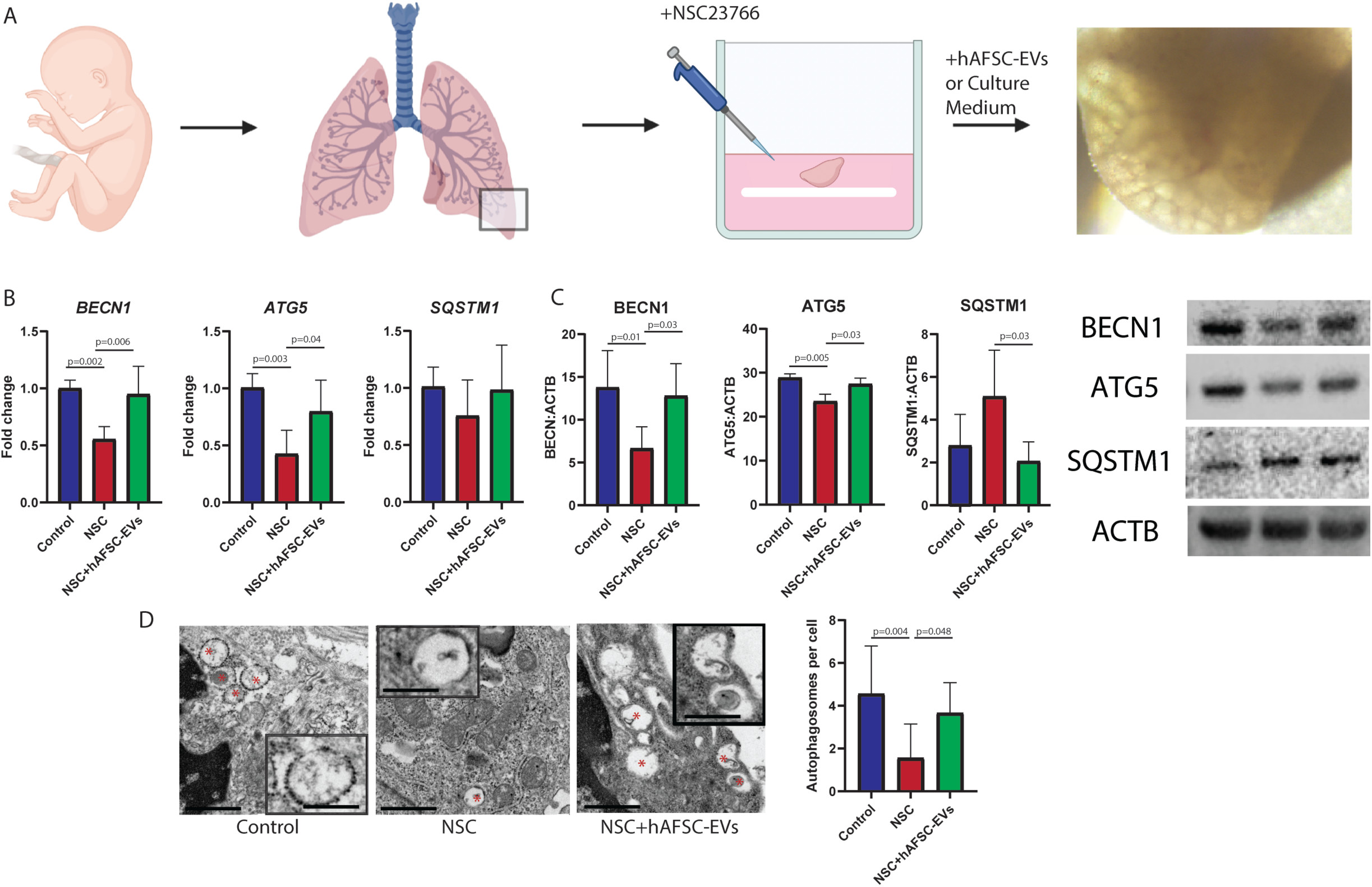
Autophagy impairment is recapitulated in a human fetal lung explant model of pulmonary hypoplasia and administration of human AFSC-EVs (hAFSC-EVs) results in autophagy restoration. (*A*) Schematic of the human fetal lung explant model based on administration of NSC23766. (*B-C*) Gene and protein expression levels of autophagy markers were impaired in explants injured with NSC23766 and restored upon treatment with hAFSC-EVs. *N*=5 lungs per group. (*D*) Fewer autophagosomes were observed in hypoplastic lung explants and autophagosome numbers were restored to normal levels following administration of hAFSC-EVs. *N*=9 cells per field per group, 4 explants per biological group. Scale bars: 1 μm (inlet scale bars: 100 nm). Data are shown as mean ± SD and were analyzed using One-way ANOVA with Tukey’s multiple comparison test. Only significant differences (p<0.05) are reported in the graphs.

## Discussion

Herein, we demonstrated that in a rat model of CDH, autophagy is reduced in the fetal hypoplastic lungs, most notably during the pseudoglandular and canalicular stages of lung development. This impairment was observed in parallel to aberrant branching morphogenesis, thus confirming that autophagy is required for proper lung development (13). Moreover, we confirmed autophagy activation in the epithelium of the developing lung, which is known to be the compartment where autophagy plays a key role in the regulation of lung homeostasis (10, 15). We replicated autophagy signaling pathway dysregulation by administering NSC23766, a RAC1 inhibitor, to a human *ex vivo* fetal lung model of pulmonary hypoplasia. Using this model, we recapitulated aberrant branching morphogenesis at pseudoglandular and canalicular stages, similar to the nitrofen-induced rat fetal model.

The mechanism behind reduced autophagy in the fetal lung remains unknown, but we speculate that in our models it could be secondary to an interaction with the retinoic acid pathway. In fact, nitrofen directly affects retinoic acid synthesis via inhibition of rate-limiting enzymes (37) and NSC23766 blocks RAC1, a downstream factor of the retinoic acid pathway (23, 38). Moreover, there is increasing evidence that retinoic acid promotes autophagy activation (39, 40). These observations are relevant to the human condition, as retinoic acid plays a role in the development of fetal lungs (41) and patients with hypoplastic lungs secondary to CDH have impaired retinoic acid status (41, 42). Moreover, fetuses with CDH have low intracellular retinoic acid levels in the lungs, as well as low retinol and retinol-binding protein levels in the cord blood (40). For this reason, several studies have tested the exogenous administration of retinoic acids as a potential strategy to promote lung growth (43-45). Although retinoic acid supplementation resulted in improvement of branching morphogenesis in the rodent model (46,47), this approach was never tested in human fetuses as retinoic acid administration during pregnancy is teratogenic (20, 48, 49).

In search for a prenatal therapy that could rescue branching morphogenesis and promote normal lung development, we recently reported the effects of an antenatal EV-based therapy. Specifically, AFSC-EVs administered to various models of pulmonary hypoplasia increased the number of branches and alveoli in fetal rat and rabbit models of pulmonary hypoplasia (5). Investigating a potential biological process influenced by AFSC-EVs, herein we reported that autophagy was rescued in both rodent and human models of pulmonary hypoplasia. AFSC-EV beneficial effects were observed at the pseudoglandular and canalicular stages, the latter of which is most clinically relevant as CDH can be diagnosed as early as this stage. The restoration of autophagy levels was observed specifically in the fetal lung epithelium that is the compartment mostly affected by the administration of AFSC-EVs (5).

When we investigated the mechanism of action of AFSC-EVs on restoration of autophagy in fetal lung tissue, we delineated the role of the miRNA cargo contained in the EVs. We identified that miR-17-5p and -20a were the most abundant members of the miR17∼92 cluster, a family of miRNAs responsible for mediating branching morphogenesis, alveolarization, and lung epithelial maturation (5). These two miRNAs have been reported to regulate autophagy through inhibition of SQSTM1 (50, 51), and our mechanistic work demonstrated that both miRNAs contributed to the ability of AFSC-EVs to restore autophagy levels. Other miRNAs that have been reported as dysregulated in hypoplastic lungs secondary to CDH (52-54) and that are contained in the AFSC-EV cargo (5), such as miR-16, -18, -19, -33, -200b, may also contribute to the AFSC-EV effects on branching morphogenesis. Given that these miRNAs do not have predicted binding sites to the mRNA of the autophagy factors of interest (BECN1, ATG5, and SQSTM1), their effect on lung morphogenesis could be through other mechanisms. Nonetheless, a major benefit of EVs is that they carry many miRNAs that are involved in regulating lung development processes, as we showed in our previous work (5).

The findings reported in this study are promising as they further explain how an AFSC-EV-based therapy could be beneficial to rescue normal fetal lung development. We recognize that several translational issues should be carefully considered before translating our findings to a clinical setting (55, 56). EVs should be tested in large animal models, such as the lamb, to address safety and feasibility, dosage, and route of administration. Towards this ultimate goal, we used GMP-grade human AFSC-EVs and confirmed similar beneficial effects in restoring autophagy in our human and rat models of pulmonary hypoplasia.

We acknowledge that this study has limitations. First, we investigated autophagy in all lung developmental stages, except the alveolar stage. This stage occurs postnatally in both rats and humans, hence not relevant for our prenatal studies. Moreover, sampling lung tissue from patients with pulmonary hypoplasia secondary to CDH after birth is considered unethical, as it would add significant morbidity to the patient. Lastly, in the mouse model, where the alveolar stage starts at the end of gestation, Yeganeh et al. showed that autophagy activity is low (13). Second, in our mechanistic experiments, we investigated only two miRNAs and it is likely that other miRNAs participate in autophagy homeostasis. We do not consider this to be a major issue, as translationally we propose to administer an EV-based rather than a miRNA-based therapy. In fact, the latter would unlikely address the multiple signaling pathways and lung compartments that are dysregulated in pulmonary hypoplasia secondary to CDH (19). Third, we used a model based on lung tissue from healthy terminations, but we recognize that the ideal platform to test human AFSC-EVs would be lung tissue from fetuses with pulmonary hypoplasia secondary to CDH. However, obtaining such specimens is challenging since fetuses with CDH terminated antenatally often have severe genetic abnormalities that do not represent typical pulmonary hypoplasia. For these reasons, to the best of our knowledge, our study is the first to employ primary fetal lung specimens at a translationally relevant timepoint for this therapy to be applied in humans.

This is the first study to show that autophagy levels are dysregulated in fetal hypoplastic lungs, thus contributing to the understanding of the pathophysiology of pulmonary hypoplasia in congenital diaphragmatic hernia. Findings from the present study corroborate the importance of autophagy in normal organ development *in utero* and could also be relevant to other congenital lung anomalies, such as bronchopulmonary sequestration and congenital pulmonary airway malformations. As autophagy plays a role in some pediatric conditions of the respiratory system, such as asthma (57), the miR-17∼92 autophagy axis herein described could also be relevant in the pathophysiology of these conditions. Finally, the present study further supports the potential of an antenatal EV-based therapy for pulmonary hypoplasia secondary to CDH. As the pathogenesis of this condition is multifactorial, an EV-based therapy containing miRNAs that address the various dysregulated biological processes, such as autophagy, would be advantageous.

## Acknowledgements

The authors would like to thank J. Rutka and C. Smith for use of NanoSight LM10, J. Reyes, D. Chiasson, and G. Somers for selection of age- and sex-matched autopsy samples, and P. De Coppi and Micregen Ltd. for providing AFSCs and AFSC-CM in kind. We are indebted to the Nanoscale Biomedical Imaging Facility, the Imaging Facility, the Lab Animal Services, and the Department of Paediatric Laboratory Medicine at the Hospital for Sick Children, Toronto, as well as the Mount Sinai Hospital Biobank, Toronto.

## Supplemental Figure Legends

**Figure E1**. Characterization of rat and human amniotic fluid stem cell extracellular vesicles. (*A*) Representative plot of the average size distribution of rat and human AFSC-EVs visualized using nanoparticle tracking analysis. Data are representative of five 40-second videos of each EV preparation. X-axis = size distribution (nm), y-axis = concentration (particles/mL). (*B*) Representative transmission electron microscopy photos of rat and human AFSC-EVs; two different magnifications highlight the morphology of individual EVs at far fields (top) and near fields (bottom). Scale bars: 1 μm for 25X images and 200 nm for 100X images. (*C*) Canonical EV markers CD63, TSG101 and Flot1 were positively detected in both rat and human AFSC-EVs and were negative for the endoplasmic reticulum marker, Calnexin (negative control marker).

**Figure E2**. RAC1 expression is decreased in the lungs of human fetuses with CDH compared to control. (*A*) Representative images of RAC1 immunofluorescence staining of the left lungs obtained from the autopsy studies of a fetus with isolated left CDH (no associated anomalies and normal karyotype) at 19 weeks of gestation and of an age- and sex-matched control (no associated anomalies, no lung pathology) Scale bars, 25 μm. (*B*) Quantification of RAC1 expression from the left lungs of two fetuses with isolated left CDH at 19 and 20 weeks of gestation, and of two age- and sex-matched controls. *N*= 24 fields per sample. Data are shown as mean ± SD and were analyzed using Student t-test.

**Figure E3**. Knockdown of miR-17-5p and miR-20a from the miR-17∼92 family results in upregulation of target genes BMPR2, HDAC4, and ATG7 in both AFSCs and AFSC-EVs. (*A-F*) *N*=3 biological replicates per group. Data are shown as mean ± SD and were analyzed using One-way ANOVA with Tukey’s multiple comparison test. Only significant differences (p<0.05) are reported in the graphs.

**Figure E4**. Branching morphogenesis is impaired by nitrofen administration and rescued by AFSC-EV treatment in rat fetal lung explants. (*A-B*) Branching was analyzed at E14.5 via brightfield microscopy to evaluate the number of terminal buds (*A*), and at E17.5 via hematoxylin and eosin staining to measure the radial airspace count (RAC; *B*). *N*=6 pups per group (*A*), *N*=3 pups per group (*B*). Scale bars, 50 μm (*A*) and 100 μm (*B*). Data are shown as mean ± SD and were analyzed using One-way ANOVA (*A*) or Kruskal-Wallis (*B*) multiple comparison tests. Only significant differences (p<0.05) are reported in the graphs.

**Figure E5**. Interrogated analysis of small RNA-sequencing data obtained from primary epithelial cells derived from hypoplastic rat fetal lungs. Compared to untreated primary epithelial cells, AFSC-EV treated cells had higher levels of miR-17 and miR-20a and decreased mRNA expression of their target SQSTM1. *N*≥ 4 per group. Data are shown as mean ± SD and were analyzed using Student t-test.

## Notes

### Competing Interest Statement

The authors have declared no competing interest.

