## Supplementary figures and images for "Autophagy is impaired in fetal hypoplastic lungs and rescued by administration of amniotic fluid stem cell extracellular vesicles"

### Supplemental Files

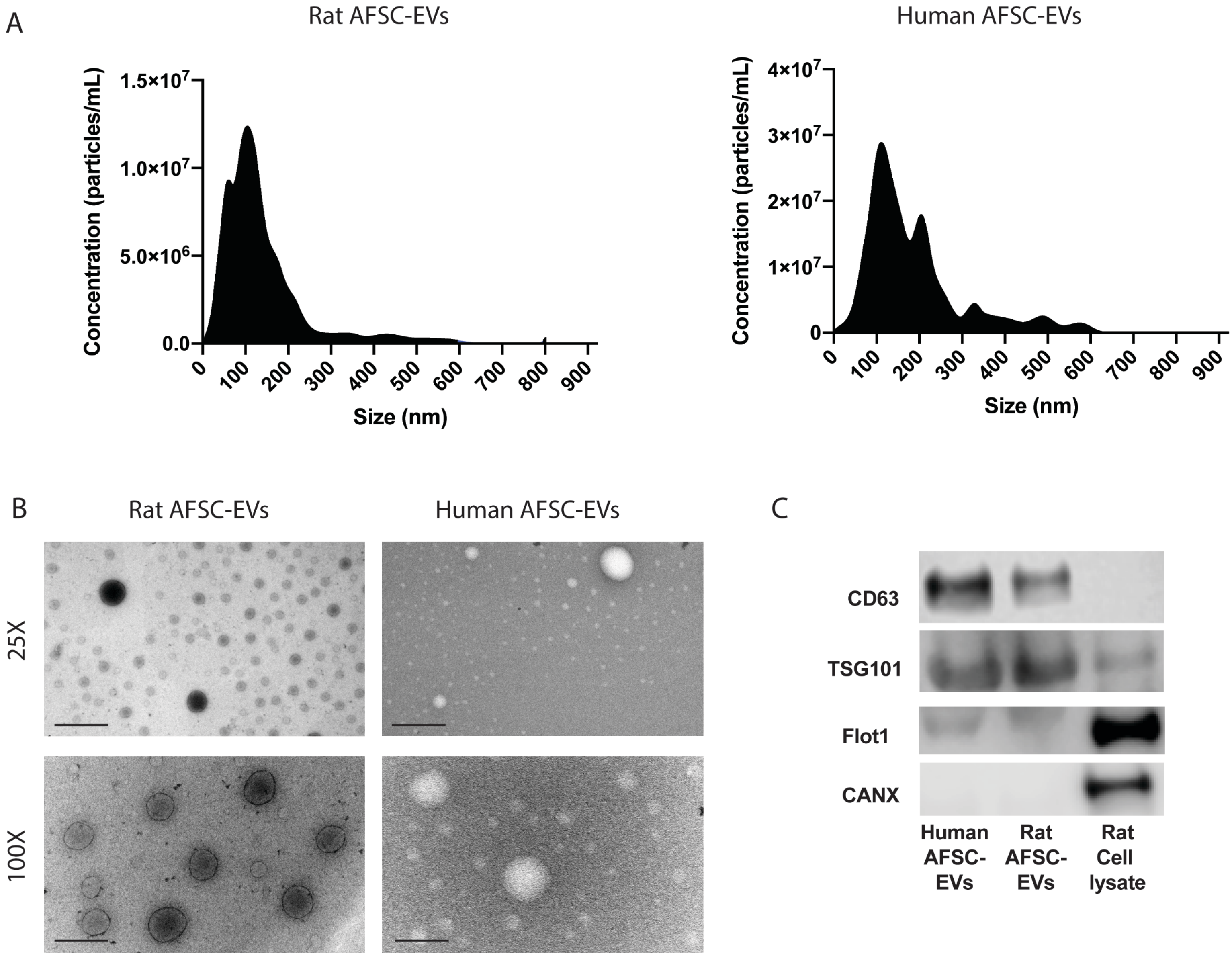

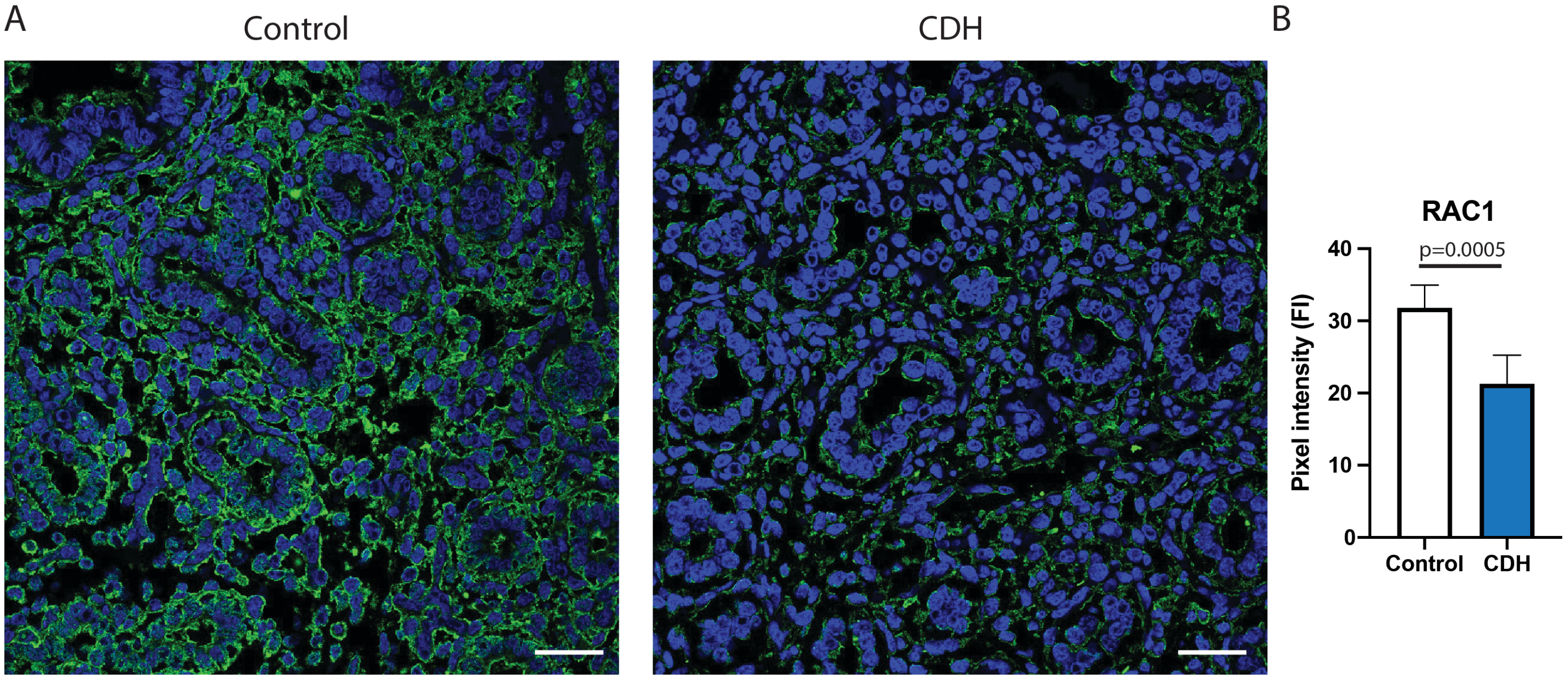

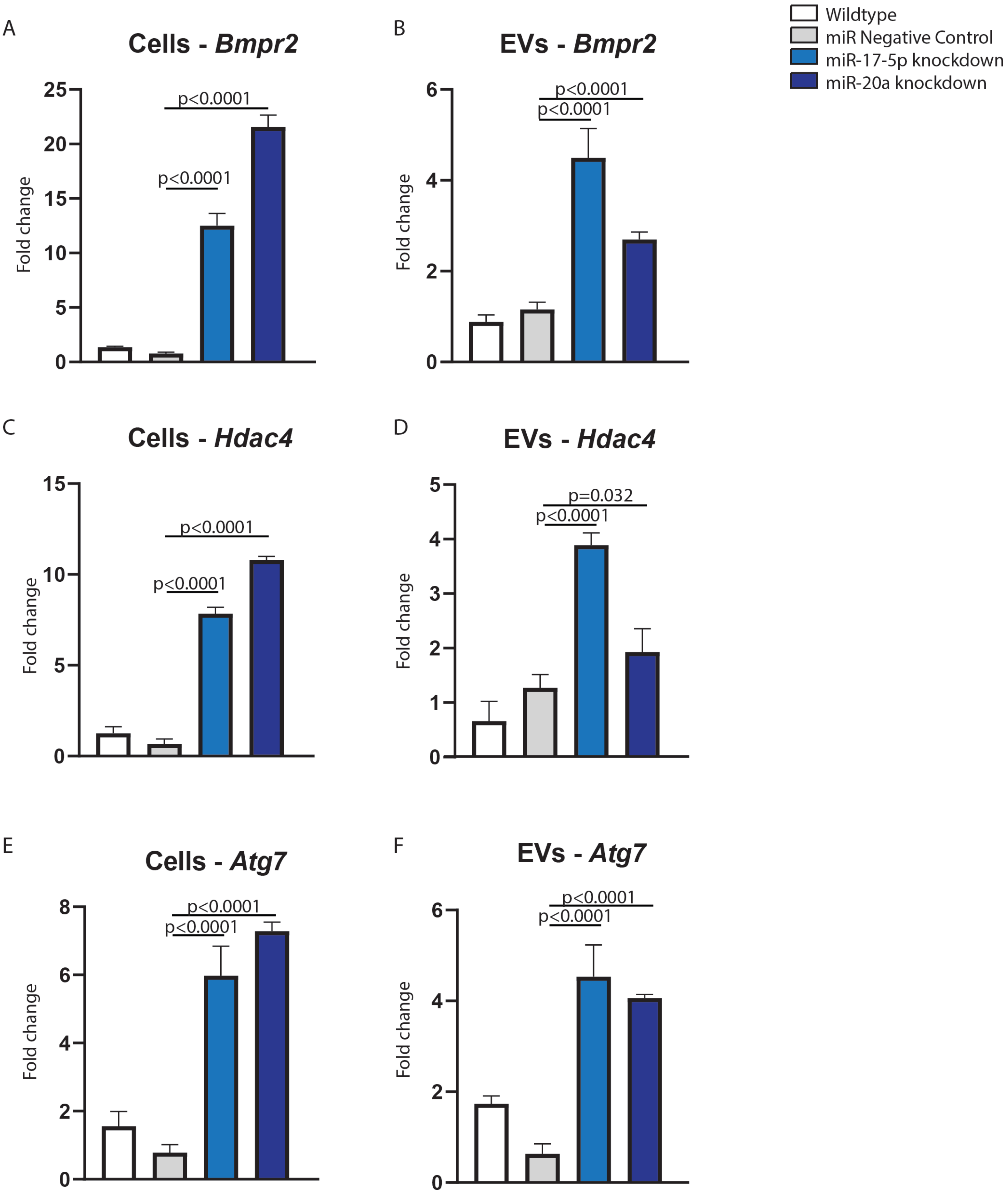

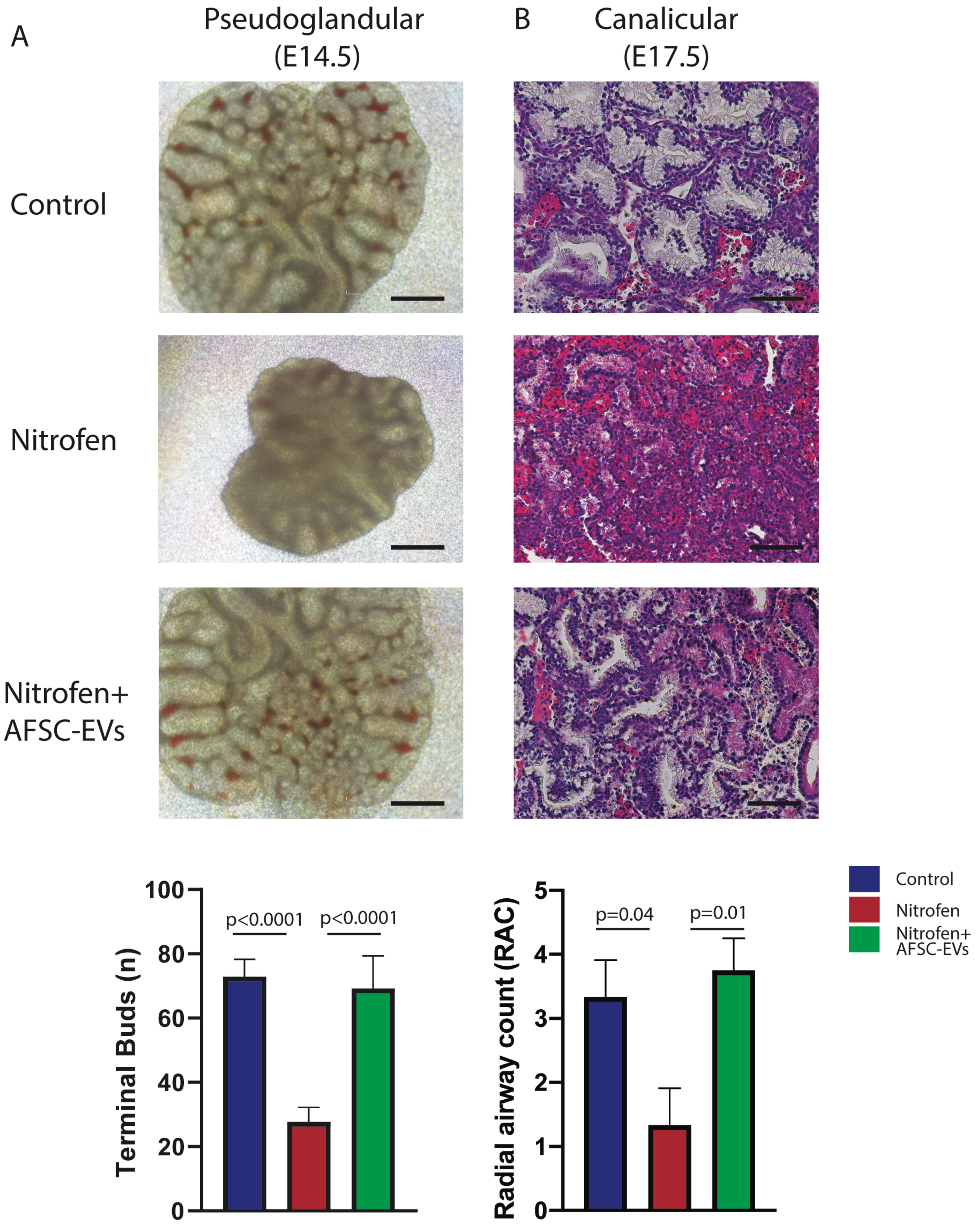

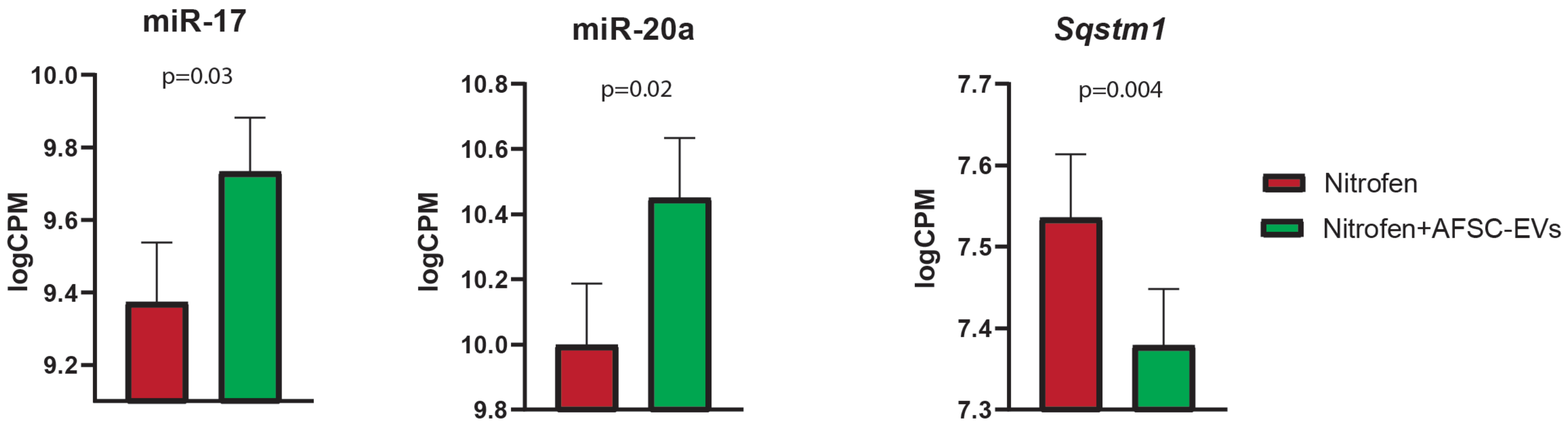
